# Transmural compression drives lymphatic endothelial dysfunction through Piezo1–CaMKII signaling

**DOI:** 10.64898/2026.11.20.768235

**Authors:** Roxane Sylvestre, Geraldine Puel, Maelle Charvoz, Perrine Singla, Tangra Draia-Nicolau, Baptiste Alric, Eric Lacazette, Anne-Catherine Prats, Yukiko Matsunaga, Barbara Garmy-Susini, Aurelien Bancaud, Florent Morfoisse

**Affiliations:** Univ Toulouse, Inserm UMR 1297, I2MC, Toulouse, France; CNRS, LAAS, Toulouse, France; Institute of Industrial Science, The University of Tokyo, 4-6-1 Komaba, Meguro-ku, Tokyo, 153-8505, Japan; Department of Bioengineering, School of Engineering, The University of Tokyo, 4-6-1 Komaba, Meguro-ku, Tokyo, 153-8505, Japan

## Abstract

**Background:** Secondary lymphedema is characterized by progressive lymphatic dysfunction, chronic inflammation, and tissue fibrosis, but how mechanical changes during disease progression contribute to lymphatic failure remains poorly understood. Lymphatic vessels initially undergo intraluminal pressure-induced dilation, whereas advanced disease is associated with extraluminal compression resulting from tissue remodeling. We investigated how these different mechanical stimuli affect lymphatic endothelial function.

**Methods and Results:** We developed a human lymphatic vessel-on-chip model enabling independent control of intra- and extraluminal pressure. In contrast to vessel dilation, extraluminal compression profoundly disrupted lymphatic endothelial barrier integrity, increasing permeability and causing junctional disorganization, while also inducing a strong pro-inflammatory transcriptional response. Mechanistically, compression induced sustained intracellular Ca²⁺ influx and activated the mechanosensitive ion channel Piezo1. Piezo1 inhibition attenuated compression-induced Ca²⁺ responses and protected endothelial barrier integrity, identifying Piezo1 as an upstream mediator of compression-induced dysfunction. Downstream, compression-induced Ca²⁺ signaling activated Ca²⁺/calmodulin-dependent protein kinase II (CaMKII), whose inhibition preserved endothelial junctional integrity and reduced both barrier dysfunction and inflammatory activation. In a murine model of secondary lymphedema, CaMKII inhibition improved lymphatic function and reduced lymphedema- associated tissue remodeling.

**Conclusions:** These findings identify a Piezo1-CaMKII signaling axis that links pathological extraluminal compression to lymphatic endothelial dysfunction and inflammation. Targeting this pathway may help preserve lymphatic function during lymphedema progression.

## Introduction

Lymphedema (LD) is a chronic and multifactorial disease that significantly reduces patients’ quality of life ^1,2^. It results from compromised lymphatic drainage leading to persistent lymph stasis, tissue swelling, and pathological tissue remodeling. LD can arise from genetic defects affecting the lymphatic vasculature (primary LD) or, more commonly in Western countries, from iatrogenic lymphatic injury following cancer treatments such as lymph node dissection, radiotherapy, or chemotherapy (secondary LD)^3,4^. The pathophysiology of LD evolves during disease progression. Following lymphatic injury, impaired lymph transport leads to the accumulation of interstitial fluid within the affected tissues. This fluid overload increases both interstitial and intraluminal hydrostatic pressures, and is associated to lymphatic vessel dilation and stretching of lymphatic endothelial cells (LECs)^5–9^. As the disease progresses, chronic lymph stasis triggers inflammation, adipose tissue expansion, excessive extracellular matrix (ECM) deposition, and progressively generates vessels narrowing and collapse^8,10^. These observations suggest that distinct forms of transmural pressure occur during LD progression: the early stages are primarily characterized by vessel dilation, whereas advanced disease is associated with compression of lymphatic vessels.

Apart from shear stress, mechanistic studies investigating the effects of mechanical forces on lymphatic vessels have remained limited. Yet, lymphatic vessels have been shown to respond to different levels of transmural pressure^6,11–13^, and LD has been associated with elevated pressure across the lymphatic wall in affected tissues^5,11,14^, highlighting the need to investigate whether alterations in this mechanical environment directly impact lymphatic vessel function. Using tissue engineering approaches^15^, we developed a lymphatic microvessel (LMV) platform that enables application of precisely controlled intraluminal (dilation) and extraluminal (compression) pressures to recapitulate mechanical cues of LD progression. We demonstrated that LMVs accommodated sustained dilation constraints with minimal alterations in vessel integrity. In contrast, compressive stress severely impaired lymphatic barrier function, increasing vessel permeability and disrupting endothelial cell-cell junctions. We then performed transcriptomic analyses of LMVs under compressive or static conditions, revealing a marked upregulation of pro-inflammatory genes in response to mechanical stimulation. These findings prompted us to investigate the molecular pathways linking mechanical stress to inflammatory activation. In this context, we identified Piezo1, a mechanically gated ion channel that acts as a key mechanosensor in LECs^16–18^, converting changes in mechanical forces into cellular signals essential for lymphatic vessel function. Its dysregulation has also been implicated in LD^19–21^, making it a critical mediator of this response. Pharmacological activation exacerbated compression-induced barrier dysfunction, whereas Piezo1 silencing protected LMVs against these deleterious effects. Among the downstream effectors of Piezo1-mediated Ca²⁺ influx, calcium/calmodulin-dependent protein kinase II (CaMKII) emerges as a prominent signaling node, coupling mechanosensitive Ca²⁺ entry to intracellular pathways that regulate endothelial function especially during inflammation^22–26^. We therefore investigated the role of CaMKII in compression-induced endothelial dysfunction and identified it as a key mediator of these deleterious effects. We found that compressive stress activated phosphorylation of CaMKII, and that pharmacological inhibition of CaMKII phosphorylation was sufficient to restore lymphatic barrier integrity and reduce inflammatory cytokine production. Using a murine model of secondary LD^27–30^, we finally proved that inhibition of CaMKII activation reduced limb swelling, improved lymphatic drainage, decreased dermal backflow and vascular leakage. Furthermore, CaMKII inhibition was associated with increased accumulation of regulatory T cells (Tregs) in lymphedematous tissues, suggesting an additional anti-inflammatory effect that may contribute to inflammation resolution. Collectively, our findings demonstrated that compressive stress is not merely a consequence of LD progression but an active driver of lymphatic dysfunction. By promoting barrier breakdown and inflammatory activation, chronic compression progressively drives the lymphatic endothelium from a functional to a dysfunctional and leaky state. Our results identify the Piezo1-CaMKII axis as a central mediator of this maladaptive response and suggest to enhance the resistance of LECs to pathological mechanical stress as a therapeutic strategy for restoring lymphatic vessel function in LD.

## Methods

### Data Availability

The data supporting the findings of this study are available from the corresponding authors upon reasonable request. RNA sequencing data are available in the National Center for Biotechnology Information Sequence Read Archive under BioProject accession PRJNA1534209.

### Animal models

All studies received local ethics review board approval and were performed in accordance with the guidelines of the European Conventionfor the Protection of Vertebrate Animals used for experimentation and according to the INSERM IACUC (France) guidelines for laboratory animal husbandry. All animal experiments were approved by the local branch Inserm Rangueil-Purpan of the Midi-Pyrénées ethics committee, France, under the reference number APAFIS ##41439-2023030811506525. Female C57BL/6J mice (6 weeks old) were obtained from Envigo, France. Animals from different cages in the same experimental group were selected to assure randomization. Mice were housed in individually ventilated cages in a temperature and light regulated room in a SPF facility and received food and water ad libitum.

### Cell culture

Human dermal lymphatic endothelial cells (HDLEC) (single male donor, juvenile foreskin, Promocell, C-12216, >99% of the cells are CD31 positive and >96% podoplanin positive) were cultured in EGM-2 MV2 medium (CC-3156, Lonza) supplemented according to the manufacturer’s instructions. Cells were maintained under standard culture conditions at 37 °C in a humidified atmosphere containing 5% CO₂, and used between passages 3 and 7 for all experiments.

### Statistical analysis

Statistical analyses were performed using GraphPad Prism software (version 10.1.2; GraphPad Software, Boston, MA, USA). For comparisons between two independent groups, unpaired two-tailed Student’s t-tests were used for normally distributed data, whereas Mann–Whitney U tests were applied for non-normally distributed data. For comparisons involving more than two groups, one-way analysis of variance (ANOVA) followed by Tukey’s multiple comparisons test was performed when parametric assumptions were met. When two independent factors were evaluated, two-way ANOVA followed by Šídák’s multiple comparisons test was used. For non-parametric comparisons involving more than two independent groups, the Kruskal–Wallis test followed by Dunn’s multiple comparisons test was applied. Data are presented as mean ± standard error of the mean (SEM), unless otherwise indicated. Statistical significance was defined as P < 0.05. Additional materials and methods are available in the Supplemental Material.

## Results

### Extraluminal compression selectively disrupts lymphatic endothelial barrier integrity

We developed a microfluidic platform to mechanically stimulate ten LMVs in parallel. Each LMV was fabricated in a silicone-based chip serving as a scaffold (see Methods) and equipped with a custom reservoir^15^, enabling the generation of a 1 cm fluid column, equivalent to 100 Pa of hydrostatic pressure. This pressure could be applied either intraluminally (causing dilation) or extraluminally (causing compression) (Figure 1A,B; Figure S1A,B). Finite element simulations (Figure 1C, Figure S1C) revealed that these changes resulted from the lymphatic barrier confining the pressure field, which created a basal-to-luminal pressure gradient that drove the deformation. The deformation of the LMV was four-fold larger in compression than in dilation (-29 ± 23% vs. + 7 ± 7%, Figure 1C), consistent with the tension–compression asymmetry of collagen gels, which exhibit elastic constants of 160 Pa and 300 Pa under −100 Pa and +100 Pa, respectively^31^. Brightfield imaging (Figure 1D, Figure S1D) experimentally confirmed these pressure-induced deformations, as reflected by the corresponding changes in vessel diameter.

**Figure 1:**
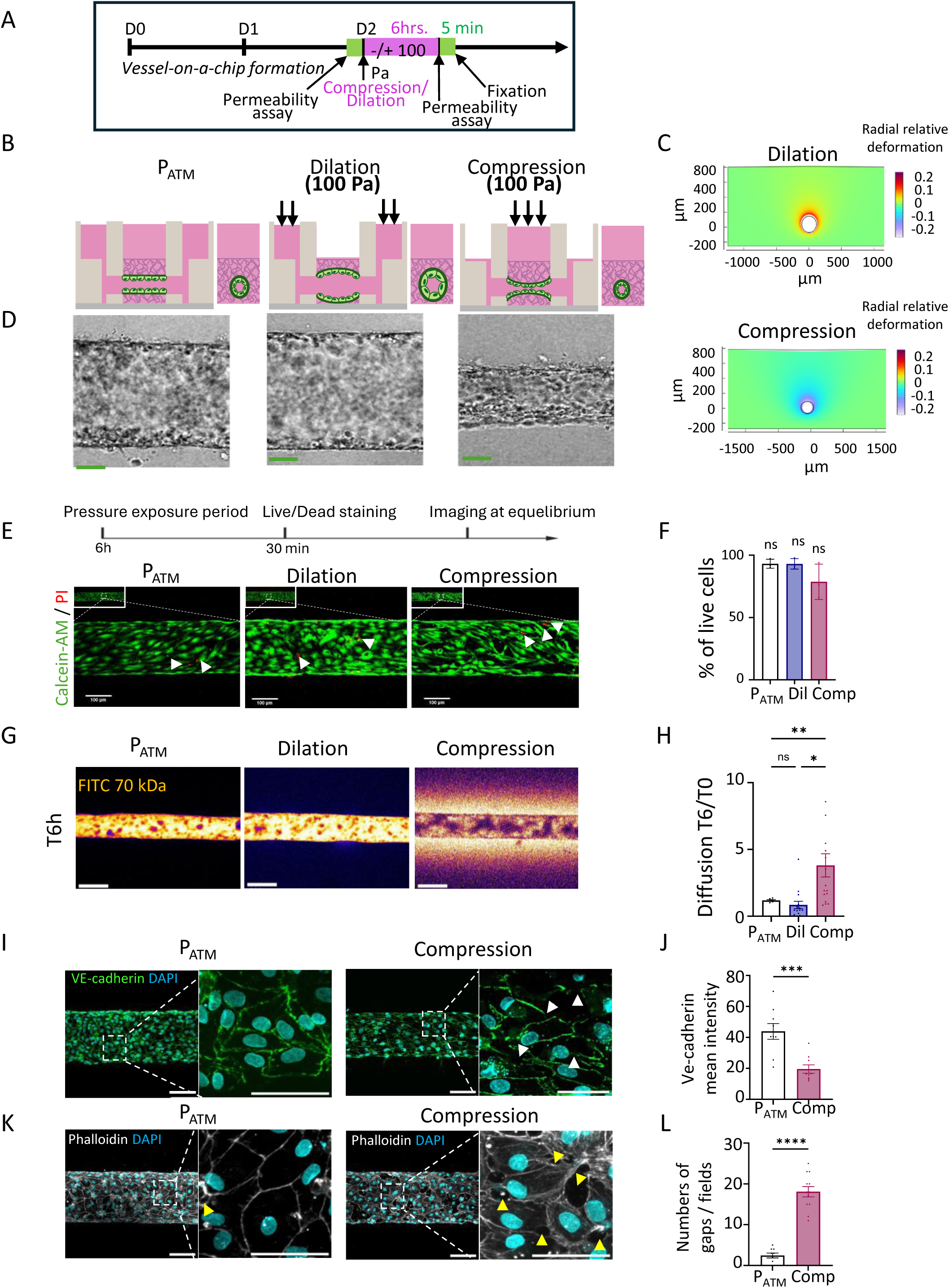
Extraluminal compression selectively disrupts lymphatic endothelial barrier integrity. **(A)** Experimental design. **(B)** Schematic representation of the 3D vessel-on-chip models composed of HDLECs embedded in type I collagen under atmospheric pressure (P_ATM_), dilation conditions (±100 Pa), or compressive conditions (-100 Pa). **(C)** Finite Element Method (FEM) simulation of the mechanical stresses applied to the chip membrane in response to a centrally applied pressure. (a) Upper panel: dilation condition (+100 Pa): radial distribution of stresses expanding outward from the region of positive pressure. Lower panel: compression condition (–100 Pa): opposite redistribution of centripetal stresses around the application point. Stress intensity is represented by a color scale (right bar), with a peak at the center. **(D)** Bright-field microscopy images of 3D HDLEC vessel-on-chip models under P_ATM_ (left panel), dilation (center) or compression (right panel). **(E)** Representative live/dead fluorescence images of vessels-on-chip under P_ATM_, dilation, and compression conditions. Calcein AM (green) labels viable cells, whereas propidium iodide (PI; red) labels non-viable cells. **(F)** Quantification of the percentage of viable cells in vessel-on-chip models under P_ATM_, dilation, and compression conditions (n = 9 per condition). Data were analyzed using one-way ANOVA. **(G)** Representative images showing fluorescent 70 kDa dextran signal intensity across the endothelial barrier at T6 h under P_ATM_, dilation, and compression conditions. **(H)** Quantification of fluorescent 70 kDa dextran diffusion coefficient into the surrounding gel, expressed as fold change (n = 12). Statistical analysis was performed using one-way ANOVA; *p < 0.05, ***p < 0.001. **(I)** Representative confocal images of VE-cadherin immunostaining highlighting endothelial cell–cell junctions (green) and DAPI-stained nuclei (blue) in vessel-on-chip models under P_ATM_, dilation, and compression conditions. Insets show higher-magnification views of selected regions. **(J)** Quantification of mean VE-cadherin signal intensity per field (n = 9). Data are presented as mean ± SD; statistical significance was determined by unpaired t-test, ***p < 0.001. **(K)** Representative confocal images of phalloidin staining highlighting the actin cytoskeleton (gray) and DAPI-stained nuclei (blue) in vessel-on-chip models under P_ATM_, dilation, and compression conditions. Insets show higher-magnification views of selected regions. **(L)** Quantification of the number of gaps per field (n = 9). Data are presented as mean ± SD; statistical significance was determined by unpaired t-test, ****p < 0.0001.

We next quantified whether compression or dilation affected endothelial barrier function by measuring the diffusive permeability^15,32,33^, which reflects the speed of diffusion from the vessel lumen to the surrounding matrix through the endothelium, using a 70-kDa FITC-dextran (Figure S1E). After 6 hr of mechanical stimulation, we measured the ratio of the final to initial diffusive permeability under static, intraluminal and extraluminal pressure conditions and observed markedly different responses across these conditions. Although neither condition induced detectable cell death (Figure 1E,F), compression, but not dilation, caused an increase in endothelial permeability, indicative of barrier degradation (Figure 1G,H, Figure S1F). Interestingly, this deleterious effect was observed only after prolonged, sustained pressure (6 hours), whereas shorter exposure times had no detectable effect on barrier function (Figure S1F,G), consistent with a previous study by Hall and colleagues showing that steady, but not pulsatile, pressure impairs LEC barrier function^34^. Consistent with these functional alterations, VE-cadherin immunostaining revealed fragmented and discontinuous adherens junctions under compression (Figure 1I,J), whereas F-actin staining showed prominent stress fiber formation and the appearance of intercellular gaps within the endothelial monolayer (Figure 1K,L), collectively indicating severe disruption of endothelial barrier integrity (Figure 1I-L). Together, these results identify sustained extraluminal compression, but not intraluminal dilation, as a potent mechanical stimulus that disrupts lymphatic endothelial barrier integrity.

## Mechanical compression triggers a pro-inflammatory transcriptional profile

To gain deeper insight into the molecular mechanisms underlying these structural and functional alterations, we performed transcriptomic analysis of LMVs exposed to compression for 6 h<u>r</u> (Figure 2A). The quality and similarity assessment of the RNA-seq samples showed distinct groups between conditions (Figure S2A, B). We then conducted a differential expression (DE) analysis, defining significantly differentially expressed genes as those with an adjusted p value (Padj) < 0.05 and a log2 fold change (log2FC) > 1 or < −1 (Figure 2B). Compression markedly affected the global transcriptome of LMVs, resulting in 1858 downregulated genes and 1222 upregulated genes, among them many genes related to endothelial activation (ICAM1, VCAM1, and SELE) and inflammation (CSF2, CSF3, CCL2) (Figure 2B and Tables S1 and S2). Notably, many known pro-inflammatory genes were among the most strongly upregulated genes in response to compression (Figure 2C in bold). Gene ontology (GO) analysis of biological processes associated with the upregulated genes confirmed this inflammatory profile, showing a significant enrichment (False Decovery Rate < 0.05) for terms related to : cytokine-cytokine receptor interaction, TNFα signaling and NFκB signaling pathway (Figure 2D). Among the upregulated genes associated with NF-κB signaling, 57 were shared between the canonical and non-canonical pathways, whereas 41 and 7 were exclusively associated with the canonical and non-canonical pathways, respectively (Figure 2E). In contrast, GO analysis of the downregulated genes revealed that no biological process was significantly enriched in compression-exposed LMVs (Figure S2C-D), although several key genes involved in the maintenance of lymphatic identity, including PROX1, LYVE1 and FLT4, were notably downregulated (Figure 2F). Overall, inflammation-related processes and responses to ions emerged as the two major functional categories among upregulated genes (Figure 2G). Taken together, these results demonstrate that extraluminal compression induces a robust pro-inflammatory transcriptional program in lymphatic vessels. Consistent with these findings, immunofluorescence analysis further revealed increased nuclear accumulation of NF-κB p65 in LMVs exposed to compression (Figure S2E).

**Figure 2:**
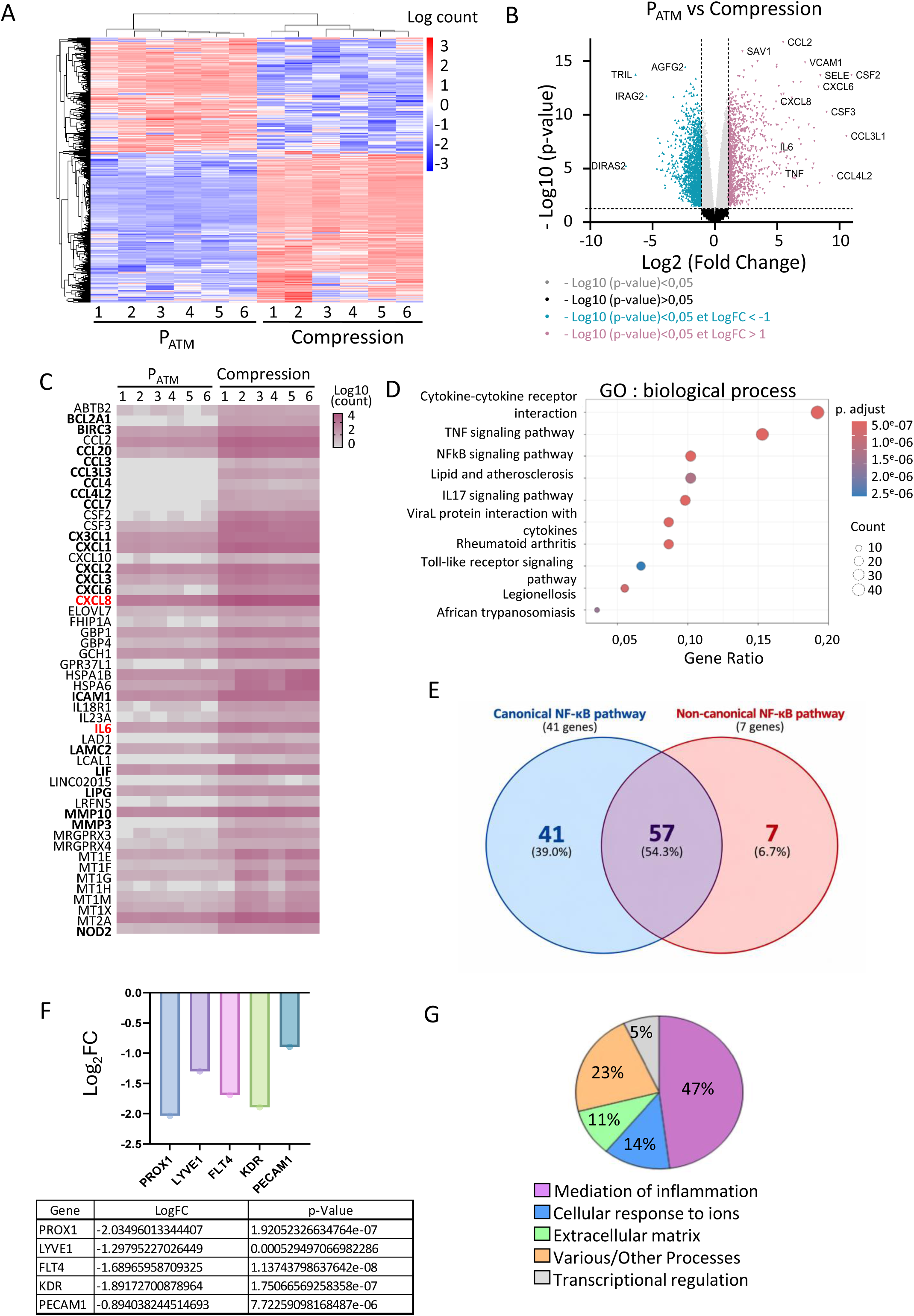
Mechanical compression triggers a pro-inflammatory transcriptional profile. **(A)** Hierarchical clustering heatmap of the differentially expressed genes across conditions. **(B)** Volcano plots showing differentially expressed genes between control vs compression. Significance threshold: adjusted p < 0.05 and |log₂ fold change| > 1. Colored dots indicate significantly upregulated (pink) or downregulated (green) genes. **(C)** Heatmap showing the expression profiles of the top 50 up-regulated genes across control and compression samples. Gene expression levels are normalized and displayed for each sample to illustrate global transcriptional changes between conditions. **(D)** Gene Ontology (GO) biological process enrichment analysis of the top 100 up-regulated genes displayed as a dot plot. Dot size represents gene count, color indicates adjusted p-value, and the x-axis shows the gene ratio. **(E)** Venn diagram showing the distribution and overlap of genes associated with the canonical and non-canonical NF-κB signaling pathways among genes significantly upregulated in compression. Percentages indicate the proportion of each gene category relative to the total number of identified genes. **(F)** Bar graph showing the log2 fold changes (Log2FC) in the expression of selected lymphatic genes. The corresponding Log2FC values and p-values are provided in the table below. **(G)** Pie chart showing the distribution of gene families among the top 100 up-regulated genes in compression.

## Piezo1 mediates compression-induced lymphatic endothelial barrier dysfunction

Given its established role as a ion-dependent mechanosensor in lymphatics^16,18^ and its involvement in mechanically induced inflammatory signaling in vascular endothelial cells^35,36^, we next investigated whether Piezo1 could mediate the response of lymphatic vessels to extraluminal compression. qPCR analysis confirmed that, among the mechanosensitive ion channels analyzed, Piezo1 was the most abundantly expressed in LECs (Figure S3A), while Fluo-8 AM imaging revealed that compression triggered a robust increase in intracellular Ca²⁺ levels in LMVs (Figure 3A,B), consistent with a potential role for Piezo1 in the response to compression. In agreement with these findings, treatment with the Piezo1 agonist Yoda1 accelerated the compression-induced calcium response, which occurred within 2 hours compared with 6 hours in control conditions (Figure S3B). Endothelial barrier disruption (Figure 3C, Figure S3C) and VE-cadherin junction fragmentation (Figure 3D) was concomitantly detected. Conversely, Piezo1 silencing using shRNA (Figure S3D) largely preserved endothelial barrier function (Figure 3E, Figure S3E) and maintained continuous cell-cell junctions despite the application of compression for 6 hours (Figure 3F). Altogether, these results establish that Piezo1 is a critical mediator of compression-induced endothelial barrier dysfunction.

**Figure 3:**
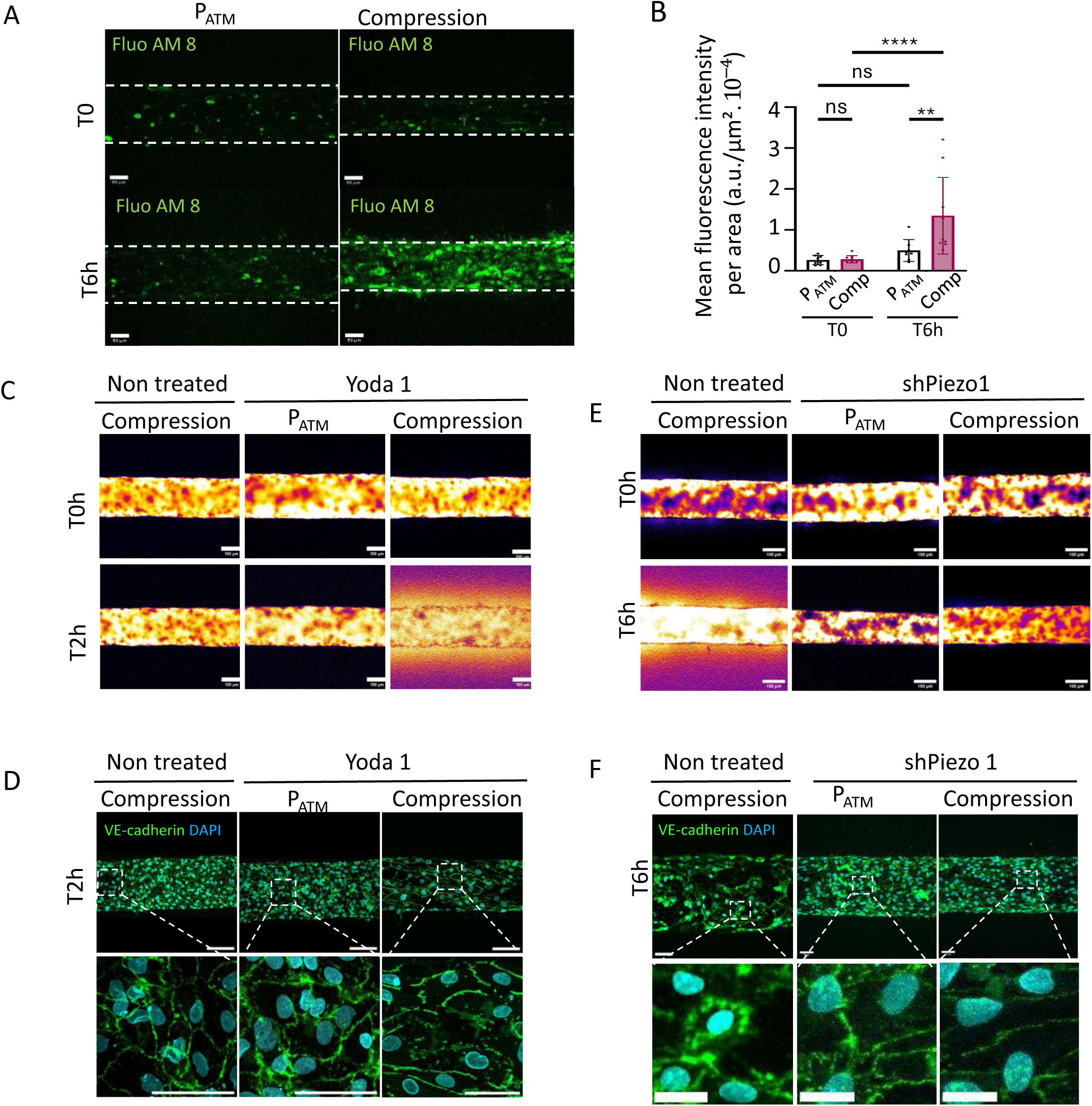
Piezo1 mediates compression-induced lymphatic endothelial barrier dysfunction. **(A)** Representative fluorescence images of Fluo-8 AM calcium dye in control and compression conditions, showing increased intracellular calcium levels under compression at 6h. **(B)** Quantification of mean fluorescence intensity per area (a.u./µm² ×10⁻⁴) is shown (n = 8 per condition), with mean ± SD. Statistical significance was determined by ANOVA (**p < 0.01, ****p < 0.0001). **(C)** Representative images showing 70-kDa fluorescent dextran across the endothelial barrier at T0 and T2 h under untreated compression, Yoda1-treated P_ATM_ and Yoda1-treated compression conditions. **(D)** Representative immunofluorescence images of VE-cadherin at endothelial cell–cell junctions under untreated compression, Yoda1-treated P_ATM_ and Yoda1-treated compression conditions. Enlarged views of the indicated regions are shown below. Nuclei are counterstained with DAPI (cyan). Scale bars, 100 µm (vessel images) and 50 µm (magnified views). **(E)** Representative images showing the fluorescence intensity of 70-kDa dextran across the endothelial barrier at T0 and T6 h under compression in untreated cells, P_ATM_ and compression conditions in Piezo1-knockdown (shPiezo1) cells. Scale bar, 100 µm. **(F)** Representative immunofluorescence images of VE-cadherin at endothelial cell–cell junctions under compression in untreated cells, P_ATM_ and compression conditions in Piezo1-knockdown (shPiezo1) cells. Enlarged views of the indicated regions are shown below. Nuclei are counterstained with DAPI (cyan). Scale bars, 100 µm (vessel images) and 50 µm (magnified views).

## CaMKII links Piezo1 activation to compression-induced endothelial dysfunction and inflammation

To further dissect the signaling cascade downstream of PIEZO1, we focused on CaMKII as a potential link between PIEZO1-mediated Ca²⁺ influx and inflammatory signaling. CaMKII is a major integrator of intracellular Ca²⁺ signals with pleiotropic roles in vascular cells^25,37^. Moreover, CaMKII has been implicated in PIEZO1-driven inflammatory signaling, including NF-κB activation, in several cellular contexts^24,26,38^. Four CAMKII isoforms have been described in mammals (α, β, γ, and δ), with α and β expression being largely restricted to the brain, whereas γ and δ are widely expressed^37,39^. As expected, only the γ and δ isoforms were detected in LECs (Figure S4A). We first demonstrated that compression gradually induced CAMKII phosphorylation (Figure 4A,B). We then evaluated the contribution of CAMKII activation on compression-induced barrier dysfunction by monitoring the permeability of LMVs treated with either KN93, the most widely used CAMKII inhibitor ^40^, or its inactive analog, KN92. We confirmed that KN93 suppressed compression-induced CAMKII phosphorylation (Figure S4B,C). Further, CAMKII inhibition by KN93 protected LMVs from the disruptive effects of compression, completely preventing the increase in vessel permeability observed in untreated and KN92-treated conditions (Figure 4C,D). Consistent with these findings, VE-cadherin and actin immunostaining revealed preserved endothelial cell-cell junctions following KN93 treatment under compression (Figure 4E,F, Figure S4D). Interestingly, Yoda1-mediated Piezo1 activation induced CaMKII phosphorylation as early as 2 hr after compression, whereas Piezo1 silencing markedly suppressed compression-induced CaMKII phosphorylation, further highlighting CaMKII as a key downstream effector of Piezo1-mediated mechanosensing (Figure S4E-H). Martin and colleagues^41^ demonstrated that CAMKII directly interacts with IKKβ, one of the IκB kinases required for the phosphorylation and release of the associated inhibitory protein IκB to promote NFκB signaling. We therefore performed a proximity ligation assay (PLA) in LMVs and found that compression markedly increased the interaction between CAMKII and IKKβ (Figure 4G,H). Accordingly, KN93 treatment abolished the compression-induced secretion of the pro-inflammatory cytokines IL6, CXCL8, and TNFα (Figure 4I), confirming a central role for CAMKII in mechanically induced inflammatory signaling. As IKKβ phosphorylation also increased following compression (Figure S4I), these findings support a model in which compression-induced CAMKII phosphorylation promotes its interaction with and phosphorylation of IKKβ, ultimately leading to NFκB activation.

**Figure 4:**
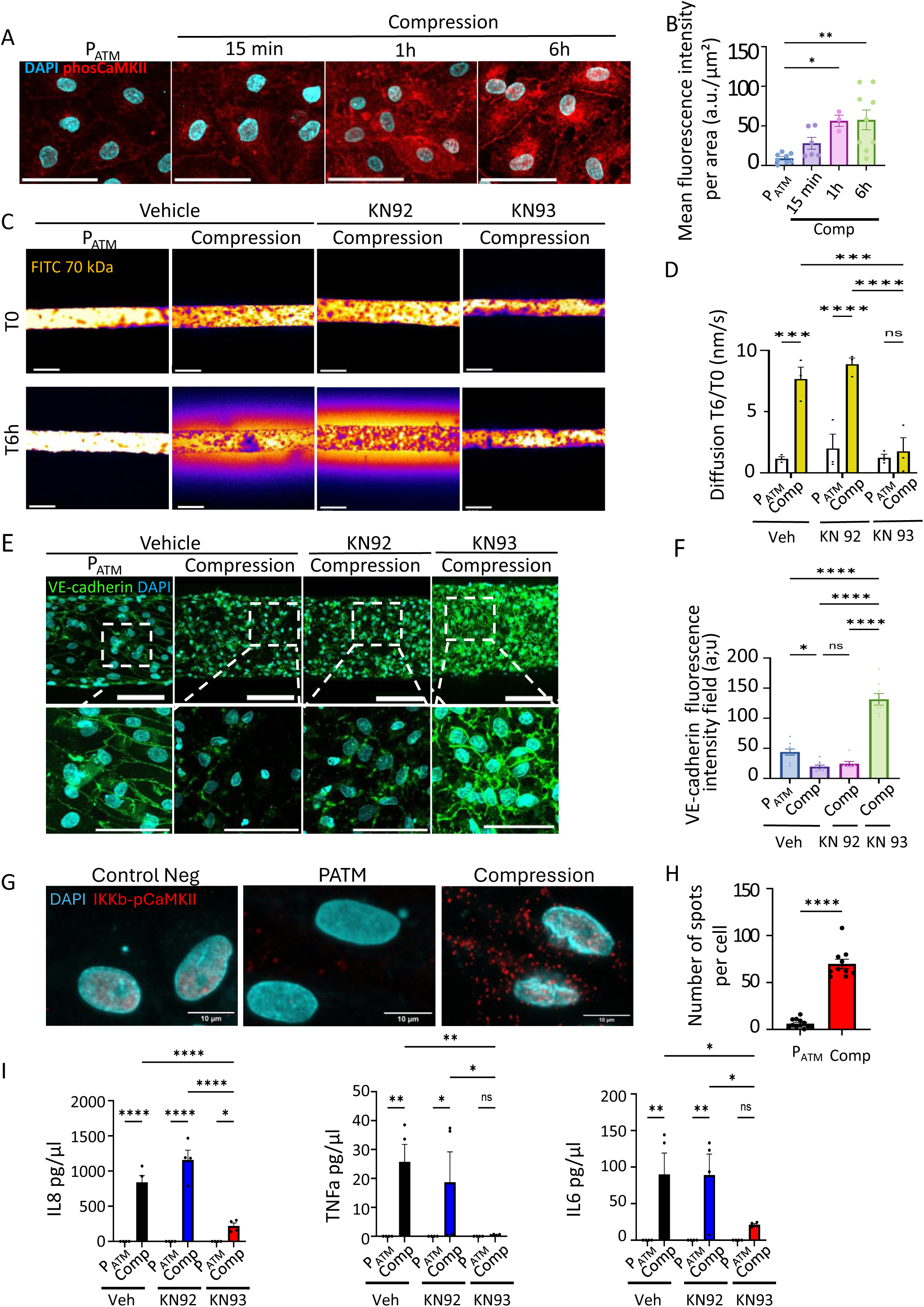
CaMKII links Piezo1 activation to compression-induced endothelial dysfunction and inflammation. **(A)** Immunostaining of phospho-CaMKII fluorescence intensity per cell in control and compressed vessels-on-chip during 15 min, 1 hr or 6 hr. Data are presented as mean ± SD (n = 9). Statistical significance was determined by unpaired t-test (**p < 0.01). **(B)** Quantification of phospho-CaMKII fluorescence intensity per cell in control and compressed vessels-on-chip during 15 min, 1 hr or 6 hr. Data are presented as mean ± SD (n = 9). Statistical significance was determined by unpaired t-test (**p < 0.01). **(C)** Representative images showing fluorescent 70 kDa dextran signal across the endothelial barrier at T0 h and T6 h under P_ATM_ and untreated, KN92-treaded and KN93-treaded compression. Scale bar: 200 µm. **(D)** Quantification of 70 kDa dextran diffusion (T6/T0, nm/s) under P_ATM_ and untreated, KN92-treaded and KN93-treaded compression. Data are presented as mean ± SD (n = 9). Statistical significance was determined by one-way ANOVA (*p < 0.05). **(E)** Representative confocal images the VE-cadherin (green) and DAPI-stained nuclei (blue) in vessel-on-chip under P_ATM_ and untreated, KN92-treaded and KN93-treaded compression. Scale bar: 100 µm. Insets show higher-magnification views of selected regions. Scale bar for insets: 20 µm. **(F)** Quantification of VE-cadherin fluorescence intensity per field (a.u.) under P_ATM_ and untreated, KN92-treaded and KN93-treaded compression. Data are presented as mean ± SD (n = 9). Statistical significance was determined by unpaired t-test (**p < 0.01). **(G)** Representative proximity ligation assay (PLA) images showing IKKβ-CaMKII interactions under negative control, P_ATM_ and compression conditions. PLA-positive signals are shown in red and nuclei with DAPI (cyan). Scale bars, 10 µm. **(H)** Quantification of PLA-positive dot per cell under P_ATM_ and compression conditions. Bars represent mean ± SEM. Statistical significance was assessed using t-test]; ns, not significant; * P < 0.05, **P < 0.01, ***P < 0.001 and ****P < 0.0001. **(I)** ELLA quantification of IL-8, TNF-α and IL-6 quantity (pg/µL) in vessel-on-chip medium maintained under P_ATM_ or compression and treated with vehicle, KN92 or KN93. Bars represent mean ± SEM. Statistical significance was assessed using ANOVA; ns, not significant; *P < 0.05, **P < 0.01, ***P < 0.001 and ****P < 0.0001.

## CAMKII inhibition preserves lymphatic function and alleviates lymphedema in vivo

As the LMVs model was designed to recapitulate the mechanical stresses sensed by LECs during LD, we next explored whether CAMKII activation could also contribute to LD progression in vivo using a murine model previously developed in the laboratory^27^. No toxicity associated with either KN92 or KN93 treatment was observed (Figure S5A–C). Mice treated with KN93 exhibited a reduced lymphedematous paw volume compared with their vehicle- and KN92-treated counterparts (Figure 5A). In control and KN92-treated mice, LD was associated with increased lymphatic vessel density, a physiopathological process known as lymphatic rerouting (Figure 5B, Figure S5D). In addition, dermal lymphatic backflow, a hallmark of dysfunctional lymphatic drainage during LD, was observed in these conditions (Figure 5B, white arrowheads, and Figure 5C). In contrast, although KN93-treated mice also displayed increased lymphatic vessel density, they exhibited markedly reduced dermal backflow, suggesting that newly formed lymphatic vessels were more functional under CAMKII inhibition (Figure 5B, white arrowheads, and Figure 5C). We further demonstrated that pCAMKII expression was increased in lymphatic vessels during LD and was abolished by KN93 treatment (Figure 5D,E). Interestingly, lymphatic vessels in the skin of LD mice treated with KN93 appeared more open and dilated compared with the other groups (Figure S5E,F) without systemic effect of either treatment in distant lymphatic networks (Figure S5G,H). Finally, Masson’s trichrome staining showed that KN93 treatment reduced LD-induced dermal fibrosis (Figure 5F,G). Taken together, these results demonstrate that CAMKII inhibition alleviates LD severity by improving lymphatic drainage capacity, limiting pathological remodeling, and reducing fibrosis.

**Figure 5:**
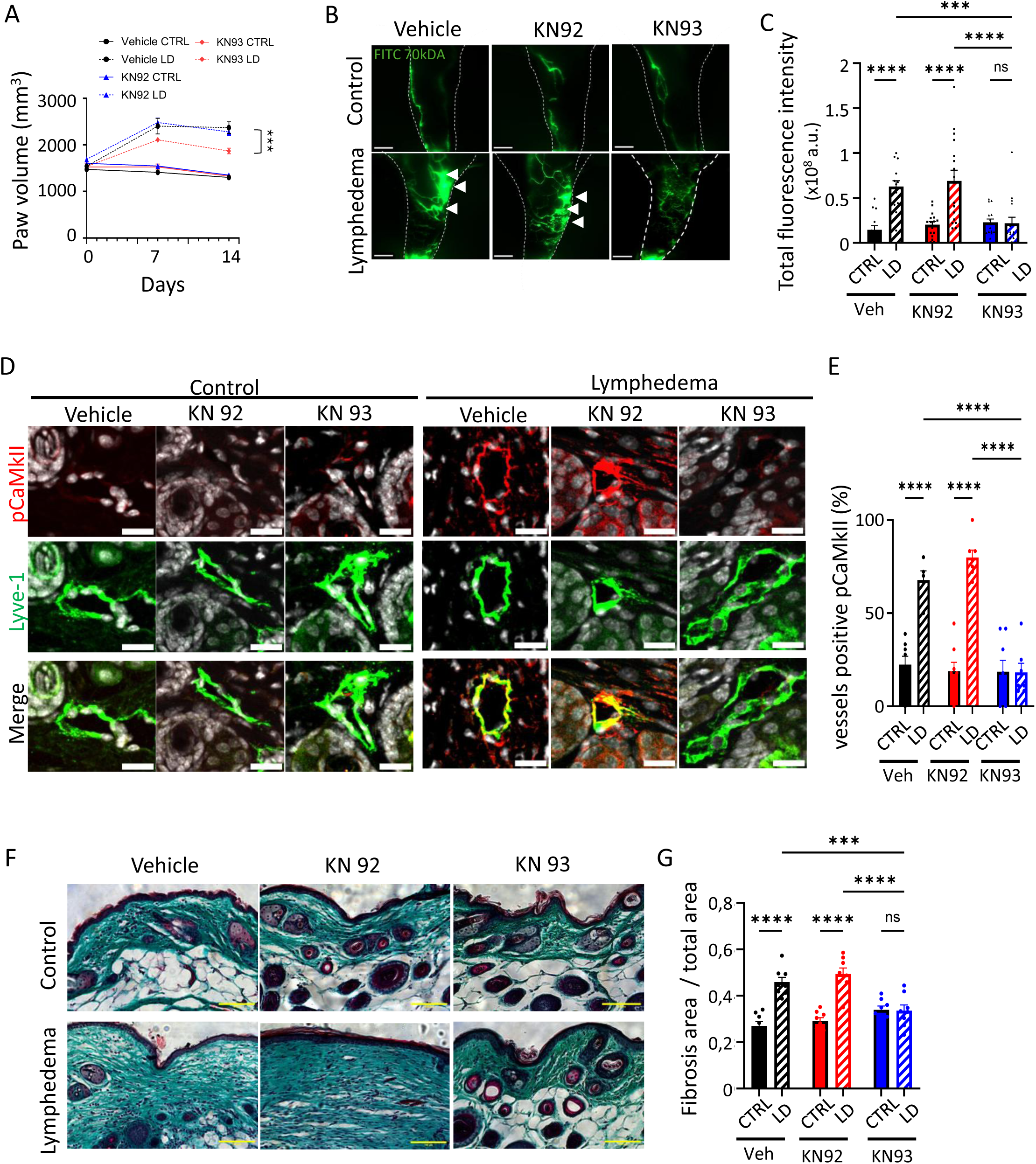
CAMKII inhibition preserves lymphatic function and alleviates lymphedema in vivo. **(A)** Paw volume measured at the indicated time points in control (CTRL) and lymphedema (LD) mice treated with vehicle, KN92 or KN93. **(B)** Representative fluorescence lymphangiography images following injection of 70-kDa FITC–dextran in control and lymphedema paws from mice treated with vehicle, KN92 or KN93. Scale bars 1 mm. **(C)** Quantification of total fluorescence intensity from FITC–dextran lymphangiography in control (CTRL) and lymphoedema (LD) paws from mice treated with vehicle, KN92 or KN93. Data are presented as mean ± SEM. n = 8 mice per condition. Statistical significance was assessed by two-way ANOVA. ns, not significant; *P < 0.05, ***P < 0.001 and *****P < 0.0001. **(D)** Representative immunofluorescence images of skin sections from control and lymphedema paws of mice treated with vehicle, KN92 or KN93, stained for the lymphatic endothelial marker LYVE-1 (green) and phosphorylated CaMKII (p-CaMKII, red). Merged images are shown in the bottom row. Scale bars 20 µm. **(E)** Quantification of the percentage of LYVE-1-positive lymphatic vessels positive for p-CaMKII in control and lymphedema paws from vehicle-, KN92- and KN93-treated mice. Data are presented as mean ± SEM. n = 8 mice per condition. Statistical significance was assessed by two-way ANOVA. ns, not significant; *P < 0.05, ***P < 0.001 and *****P < 0.0001. **(F)** Representative images of Masson’s trichrome-stained skin sections from control and lymphedema paws of mice treated with vehicle, KN92 or KN93, showing tissue fibrosis. Scale bars 100 µm. **(G)** Quantification of fibrosis, expressed as fibrotic area relative to the total tissue area, in control and lymphedema paws from vehicle-, KN92- and KN93-treated mice. Data are presented as mean ± SEM. n = 8 mice per condition. Statistical significance was assessed by two-way ANOVA. ns, not significant; *P < 0.05, ***P < 0.001 and *****P < 0.0001.

## CAMKII inhibition promotes Treg accumulation in lymphedematous tissue

Beyond improving lymphatic function, our on-chip experiments identified an anti-inflammatory effect of CAMKII inhibition. In LD, regulatory T cells (Tregs) have been implicated in the control of tissue inflammation and the resolution of chronic inflammatory responses^28,42^. We therefore investigated whether the beneficial effects of KN93 treatment *in vivo* were associated with changes in the local T lymphocytes population using flow cytometry analysis. As previously shown^30^, no effect of CD8^+^ T cell population was found in LD (Figure 6A,B) whereas CD4^+^ T cells increased in lymphedematous tissues from KN93-treated mice (Figure 6C). Among these lymphocytes, we next studied FoxP3^+^ Tregs revealing an increased Treg abundance in lymphedematous tissues from KN93-treated mice (Figure 6D,E), whereas no significant increase was observed in vehicle- or KN92-treated mice. Immunofluorescence analysis further confirmed increased Treg accumulation in the skin of KN93-treated mice (Figure 6F,G) during LD. These findings revealed that CAMKII inhibition, not only preserves lymphatic function but may also promote inflammation resolution in LD further supporting CAMKII as a potential therapeutic target.

**Figure 6:**
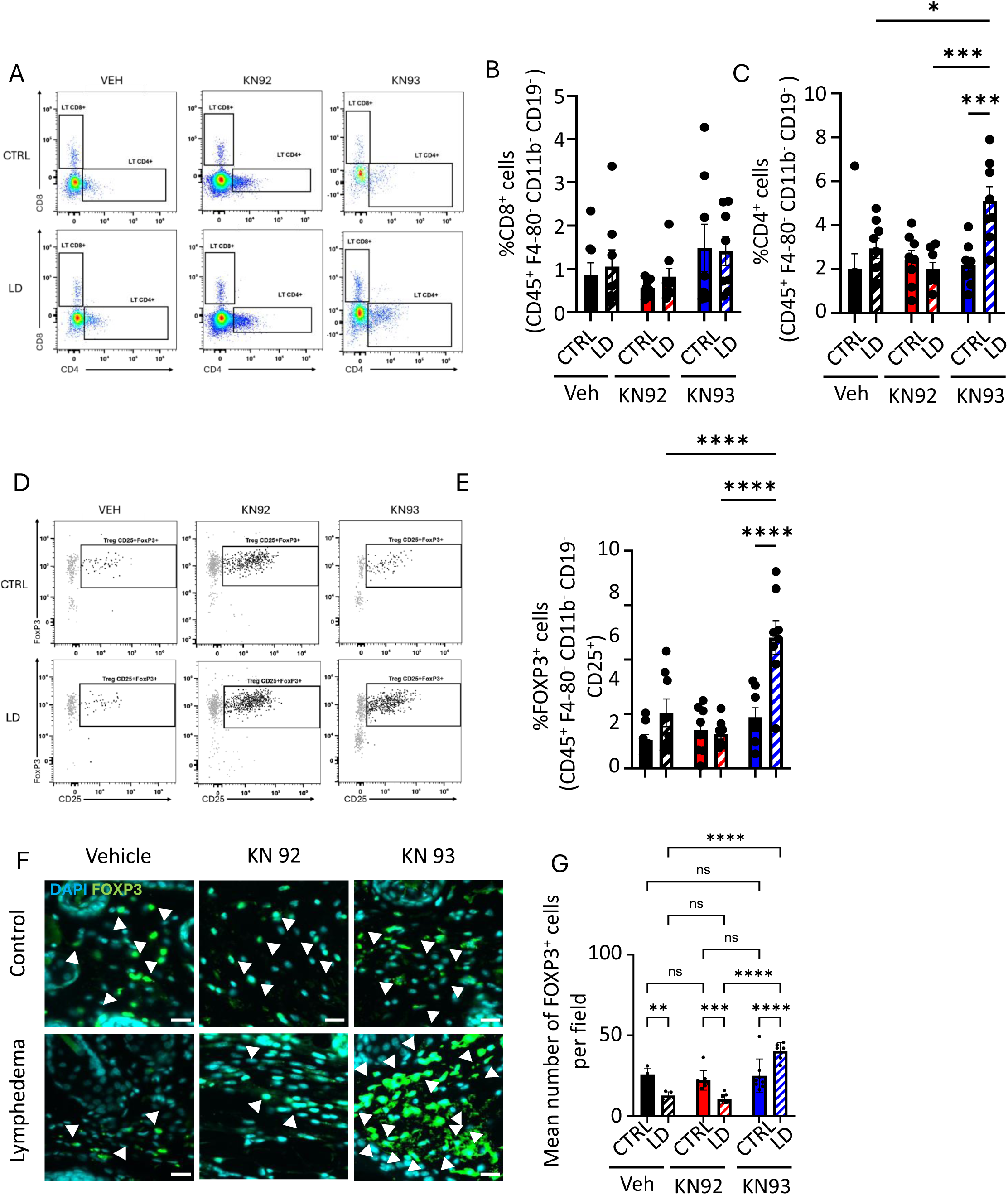
CAMKII inhibition promotes Treg accumulation in lymphedematous tissue (A) Representative FACS plots of CD8^+^ and CD4^+^ T lymphocytes in the CTRL and LD skins from mice treated with vehicle, KN92 or KN93. (B) Quantification showing the proportion of CD8^+^ T cell population in each group. (C) Quantification showing the proportion of CD4^+^ T cell population in each group. (D) Representative FACS plots of FoxP3^+^ T lymphocytes in the CTRL and LD skins from mice treated with vehicle, KN92 or KN93. (E) Quantification showing the proportion of FoxP3^+^ cell population in each group. Data are presented as mean ± SEM. n = 8 mice per condition. Statistical significance was assessed by two-way ANOVA. ns, not significant; *P < 0.05, ***P < 0.001 and *****P < 0.0001. (F) Representative immunofluorescence images of skin sections from CTRL and LD paws of mice treated with vehicle, KN92 or KN93, stained for the Treg lymphocytes marker FoxP3 (green). Scale bars 20 µm. (G) Quantification of the number of FoxP3^+^ Treg lymphocytes in control and lymphedema paws from vehicle, KN92- and KN93-treated mice. Data are presented as mean ± SEM. n = 8 mice per condition. Statistical significance was assessed by two-way ANOVA. ns, not significant; *P < 0.05, ***P < 0.001 and *****P < 0.0001.

## Discussion

Mechanical forces are increasingly recognized as key regulators of lymphatic vessel biology, controlling lymphatic valve development, vessel maturation, contractile function, and the maintenance of lymphatic endothelial identity^43,44^. However, the impact of transmural pressure, particularly under pathological conditions such as LD, has remained largely underexplored. This knowledge gap largely reflects the lack of experimental models capable of isolating pressure-induced mechanical cues from the numerous biochemical changes that occur simultaneously during disease progression in vivo. Here, we address this limitation using engineered LMVs that enable the independent application of intraluminal or extraluminal pressure, thereby reproducing the two successive mechanical states experienced by lymphatic vessels during LD progression: dilation and compression.

First, we demonstrate that LECs respond fundamentally differently to dilation and compression. In accordance with previous studies performed on blood vessel on-chip^32,33^, sustained intraluminal pressure was remarkably well tolerated. At the opposite, chronic exposure to extraluminal compression profoundly weakened cell-cell junctions by inducing VE-cadherin disorganization, altering endothelial barrier function thus increasing vascular permeability and leakiness. promoted inflammatory activation, and impaired lymphatic function recapitulating features characteristic of dysfunctional lymphatic vessels observed in LD. Transcriptomic profiling further demonstrated that compression induces a strong endothelial activation signature characterized by inflammatory cytokine production (CSF2, CSF3), adhesion molecule expression (ICAM1, VCAM1), and upregulation of canonical NFκB targets (IL6, CXCL8, TNFα), while key genes involved in the maintenance of lymphatic endothelial identity (PROX1, VEGFR3) were notably downregulated. Rather than representing a passive consequence of pathological progression, mechanical compression emerges as an active driver of disease amplification capable of reshaping LEC phenotype shifting from a homeostatic lymphatic endothelial state toward a pro-inflammatory and dysfunctional phenotype further accelerating lymphatic dysfunction. More broadly, these findings identify lymphatic mechanobiology as a fundamental yet largely overlooked component of LD pathophysiology and underscore the importance of understanding how evolving mechanical cues regulate LECs fate during disease progression.

A central question raised by these findings is how LECs sense and translate extraluminal compression into a pro-inflammatory response. Mechanosensitive ion channels are attractive candidates for this process, as they provide a direct molecular interface between mechanical forces and intracellular signaling. Here, we identify Piezo1 as a critical mediator of compression-induced endothelial dysfunction: while pharmacological activation of Piezo1 markedly accelerated barrier disruption in response to compression, genetic silencing of Piezo1 preserved endothelial integrity despite sustained exposure to compression, demonstrating that Piezo1 is functionally required to initiate the pathological response. Importantly, these findings should not be interpreted as indicating that Piezo1 activation is inherently detrimental. On the contrary, Piezo1 is required for normal lymphatic physiology. Previous studies have established its essential roles in lymphatic development, valve morphogenesis, maintenance of vessel architecture, and adaptation to fluid shear stress^16–18,44^. Rather, our results highlight the context-dependent nature of mechanotransduction, whereby a sensor that is essential for physiological adaptation becomes maladaptive when exposed to chronic pathological mechanical stimulation. This concept is further supported by human genetics, as both gain- and loss-of-function mutations in PIEZO1 cause generalized lymphatic dysplasia^19–21^, underscoring that lymphatic homeostasis relies on the precise tuning of Piezo1 activity.

Calcium influx represents the primary consequence of Piezo1 activation, yet the downstream signaling events are numerous, therefore identifying the pathway responsible for the compression-induced endothelial dysfunction is a critical point. We demonstrated that PIEZO1- mediated Ca²⁺ influx activates downstream CAMKII signaling, which can serve as a critical link between mechanotransduction and inflammatory pathways. In particular, CaMKII activation can promote NF-κB signaling^26,38,45^, thereby coupling mechanical stress to the induction of pro-inflammatory gene expression in LECs. Following compression, CaMKII phosphorylation was increased, and inhibition of CaMKII completely prevented barrier disruption, loss of VE-cadherin and preserve cell-cell junction integrity. Previous work demonstrated that CaMKII can interact with IKKβ^41^, one of the IκB kinases regulating NFκB activation. Consistent with this mechanism, we observed increased CaMKII–IKKβ interaction and enhanced IKKβ phosphorylation following compression, both of which were inhibited by KN93 treatment. In addition, inhibition of CaMKII prevented compression-induced inflammatory cytokine secretion. As a kinase, our findings strongly suggest that CaMKII phosphorylates IKKβ in response to compression, thereby promoting the phosphorylation of the inhibitory protein IκB and its subsequent dissociation from NF-κB, ultimately triggering NF-κB activation.

The therapeutic relevance of this pathway was demonstrated in vivo using a mouse model of secondary LD where we confirmed an increase phosphorylation of CaMKII in lymphatic vessel during LD. Accordingly, pharmacological inhibition of CaMKII significantly reduced LD volume, and LD-associated fibrosis. Interestingly, CaMKII inhibition did not prevent LD- associated lymphangiogenesis, but rather improved the functionality of newly formed lymphatic vessels, as indicated by decreased dermal backflow and vascular leakage reflecting an improved lymphatic drainage. This distinction is important as numerous studies have demonstrated that increased lymphatic vessel density does not necessarily correlate with efficient lymphatic transport^29,46,47^. Together, these findings suggest that preserving the functional competence of the lymphatic endothelium through CaMKII inhibition may reduce disease severity by limiting the transition from the early and reversible stages of LD to the late stages, where self-sustaining fibrosis, tissue thickening, and lymphatic vessel compression drive irreversible disease progression.

In conclusion, this study identifies extraluminal compression as a previously underappreciated driver of lymphatic endothelial dysfunction and establishes mechanotransduction as a central determinant of LD progression. We reveal a mechanistic pathway linking pathological mechanical stress to sustained inflammation through Piezo1-dependent calcium signaling and CaMKII activation. By identifying CaMKII as a pharmacologically actionable signaling node capable of restoring lymphatic vessel function and limiting fibrosis in vivo, our work expands the current understanding of lymphatic failure within the fibrotic microenvironment and opens new therapeutic perspectives aimed not only at reducing inflammation but also at enhancing the mechanical resilience of lymphatic vessels to their pathological environment.

## Acknowledgments

We thank C. Segura, and especially R.Flores-Flores for the outstanding technical assistance (GenoToul PHY platform). We thank A. Zakaroff, E. Riant (TRI platform), C.Bernis,M.Lemarinier and A.Lucas (We-Met platform) for the technical support. We thank the platform Anexplo Genotoul- CREFRE (Inserm US006, Toulouse, France) for their technical advices regarding mice experiments. We thank the Zellidja Foundation for the Travel Grant of Roxane Sylvestre.

## Sources of Funding

This study was partially supported by the grant EUR (Ecole Universitaire de Recherches) CARe N°ANR-18-EURE-0003 in the framework of the “Programme des Investissements d’Avenir”. We thank the CARe Graduate School for the support of Roxane Sylvestre.

## Disclosures

The authors declare no conflicts of interest.

## Notes

### Competing Interest Statement

The authors have declared no competing interest.

